# Defining unique structural features in the MAFA and MAFB transcription factors that control *Insulin* gene activity

**DOI:** 10.1101/2023.08.23.554429

**Authors:** Jeeyeon Cha, Xin Tong, Katie C. Coate, Min Guo, Jin-hua Liu, Garrett Reynolds, Emily M. Walker, Richard A. Stein, Hassane Mchaourab, Roland Stein

**Author notes:** Co-first authors (Jeeyeon Cha, Xin Tong, Katie C. Coate).

## Abstract

MAFA and MAFB are related basic-leucine-zipper domain containing transcription factors which have important overlapping and distinct regulatory roles in a variety of cellular contexts, including hormone production in pancreatic islet α and β cells. Here we first examined how mutating conserved MAF protein-DNA contacts obtained from X-ray crystal structure analysis impacted their DNA-binding and *Insulin* enhancer-driven activity. While most of these interactions were essential and their disruption severely compromised activity, we identified that regions outside of the contact areas also contributed to activity. AlphaFold 2, an artificial intelligence-based structural prediction program, was next used to determine if there were also differences in the three-dimensional organization of the non-DNA binding/dimerization sequences of MAFA and MAFB. This analysis was conducted on the wildtype (WT) proteins as well as the pathogenic MAFA^Ser64Phe^ and MAFB^Ser70Ala^ *trans*-activation domain mutants, with differences revealed between MAFA^WT^ and MAFB^WT^ as well as between MAFA^Ser64Phe^ and MAFA^WT^, but not between MAFB^Ser70Ala^ and MAFB^WT^. Moreover, dissimilarities between these proteins were also observed in their ability to cooperatively stimulate *Insulin* enhancer-driven activity in the presence of other islet-enriched transcription factors. Analysis of MAFA and MAFB chimeras disclosed that these properties were greatly influenced by unique C-terminal region structural differences predicted by AlphaFold 2. Importantly, these results have revealed features of these closely related proteins that are functionally significant in islet biology.

## INTRODUCTION

The musculoaponeurotic fibrosarcoma (MAF) family consists of proto-oncogenes with important cellular regulatory properties that are categorized into two distinct subgroups: small MAFs that lack an N-terminal *trans*-activation domain (∼150-160 amino acids: MAFF, MAFG, and MAFK) and large MAFs (∼240-340 amino acids: MAFA, MAFB, c-MAF, and NRL) (Artner et al., 2010; Conrad et al., 2015). These MAF proteins act as transcription factors (TFs) and play active roles in many tissue types, such as the pancreas (Matsuoka et al., 2003; Sakai et al., 1997), lens (Sakai et al., 1997), and cartilage (MacLean et al., 2003) by contributing to their development, differentiation and regulation of mature function.

The MAFA and MAFB proteins have been found to be vital in the development and/or maturation of pancreatic islet insulin hormone-producing β- and glucagon hormone-producing α-cells (Artner et al., 2010; Conrad et al., 2015). Expression of these proteins is affected in type 2 diabetes, a disease ultimately arising from insulin deficiency and concomitant glucose under-utilization, which is compounded by glucagon excess driving hepatic glucose over-production (Guo et al., 2013). Moreover, mutations near a priming kinase site in both MAFA and MAFB are clinically relevant. The Ser64Phe (i.e., S64F) substitution prevents S65 phosphorylation of MAFA and produces either monogenic diabetes or insulinomatosis (Iacovazzo et al., 2018), while the Ser70Ala (i.e., S70A) mutation in MAFB blocks S70 phosphorylation and causes multicentric carpotarsal osteolysis (i.e., MCTO), a pediatric multi-system disorder characterized by osteolysis of the carpal and tarsal bones, subtle craniofacial deformities, and nephropathy (Ma et al., 2023). While MAFB appears to be essential to insulin production during human β cell development (Russell et al., 2020), it is unclear whether MAFB^S70A^ can cause diabetes in MCTO patients.

The X-ray crystal structure has been determined for the basic-leucine-zipper regions of MAFA (Lu et al., 2012a) and MAFB (Textor et al., 2007), which are responsible for DNA binding and homo- or hetero-dimerization (Blank and Andrews, 1997). Here we first analyzed how mutations in conserved basic region amino acids which interacted with DNA control element sequences in these crystallographic studies influenced *Insulin* enhancer-driven and DNA-binding activity of the full-length protein. While most of these interactions were critical, one variant of MAFA (Arg272Ala, or MAFA^R272A^) functioned similarly to the wildtype (WT) protein while its analogous mutation in MAFB was debilitating, suggesting contributions from regions lying outside of the basic-leucine-zipper domains. To further analyze these areas, we applied AlphaFold 2 algorithm to obtain insight into their underdefined N-terminal *trans*-activation domain and C-terminal region structure. Functional and molecular assays were employed to evaluate the relevance of the predicted structural differences to the activity of MAFA^WT^, MAFB^WT^ and their pathogenic variants. Our results strongly suggest that structural dissimilarities between MAFA and MAFB impart their unique functional properties under physiologic and disease conditions.

## RESULTS

### 1) In contrast to MAFB^R256A^, the analogous MAFA^R272A^ mutant does not impact *trans*-activation or DNA binding activity

The amino acids comprising the basic region of MAFA and MAFB are absolutely conserved as were *cis*-control element DNA contacts within the *Escherichia coli* expressed basic leucine-zipper portion of the protein (i.e., MAFA, residues 226–318; MAFB, 211–323) analyzed previously by X-ray crystallography (**Figure 1A**) (Lu et al., 2012a; Textor et al., 2007). We first determined how mutation of four amino acids within their basic region would affect the *trans*-stimulation and DNA-binding activity of full-length MAFA and MAFB, considering the possibility that post-translational events (e.g., phosphorylation (Guo et al., 2010), ubiquitination (Guo et al., 2009), and sumoylation (Shao and Cobb, 2009)) produced within these heavily modified proteins in mammalian cells could be influential. Of note, the X-ray crystal structure revealed that three of the analyzed residues in MAFA (i.e., R260, R265, and Y267) and MAFB (R244, R249, Y251) have essentially the same rotameric alignment in relation to the *cis*-acting control element DNA sequences and have near-complete overlap in chains A and B of the protein homodimer (**Figure 1B**). In contrast, even though MAFA^R272^ and MAFB^R256^ contact the same control element guanine, they utilized a distinct rotameric state for binding (Lu et al., 2012b; Pogenberg et al., 2014) (**Supp Figure 1**).

**Figure 1.**
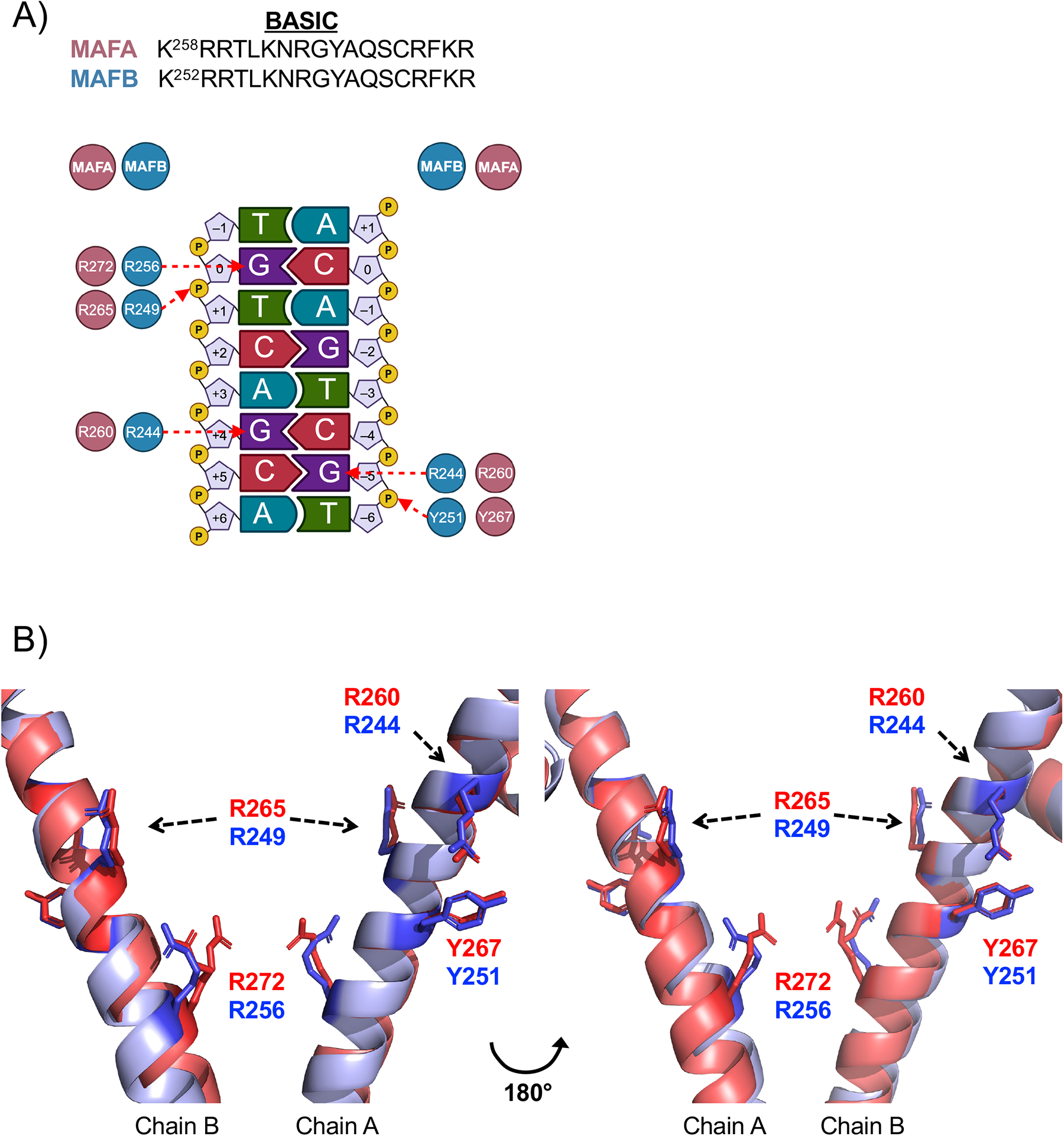
X-ray crystal structure analysis revealed that conserved basic region-DNA binding MAFA^R272^ has a unique rotameric structure to the analogous MAFB^R256^. A) Top: The sequences of basic region are identical between MAFA, MAFB, and the other closely related large MAF proteins (i.e., c-MAF and NRL (Lu et al., 2012a). Bottom: Schematic representation of X-ray crystallographic results illustrating the MAFA and MAFB basic region amino acid base-specific interactions under analysis. B) Opposite views of the DNA overlay of the crystal structures for MAFA (4EOT, red) (Pogenberg et al., 2014) and MAFB (2WTY, blue) (Lu et al., 2012b). The four residues shown in stick schematics are R260, R265, Y267, and R272 of MAFA and R244, R249, Y251, and R256 of MAFB.

The three conserved arginines (i.e., R) and the unique tyrosine (Y) of the MAFA and MAFB basic region were mutated to alanine (A) and phenylalanine (P), respectively. The full length WT and mutant proteins were produced in transfected HeLa cells and analyzed for their ability to stimulate *Insulin*-enhancer driven reporter activity, bind *Insulin* control element DNA in gel shift assays, and stably produce protein by immunoblotting. All of the mutants were expressed at similar levels (**Supp Figure 2**), and each of the basic region DNA-binding mediating mutants of MAFB (i.e. R244A, R249A, Y251F, R256A) and most in MAFA (R260A, R265A, Y267A/F/S; **Figure 2A**) were debilitating in the reporter assays. However, MAFA^R272A^ had WT-like activity whereas the comparable MAFB^R256A^ mutant was inactive (**Figure 2A**). Notably, the MafA antibody super-shifted MAFA^R272A^ and competed specifically in gel shift assays, whereas the dysfunctional mutants did not (**Figure 2B**, data not shown). The results indicated that structural features in MAFA and MAFB residing outside the region analyzed earlier (Lu et al., 2012a; Textor et al., 2007) were consequential to activity.

**Figure 2.**
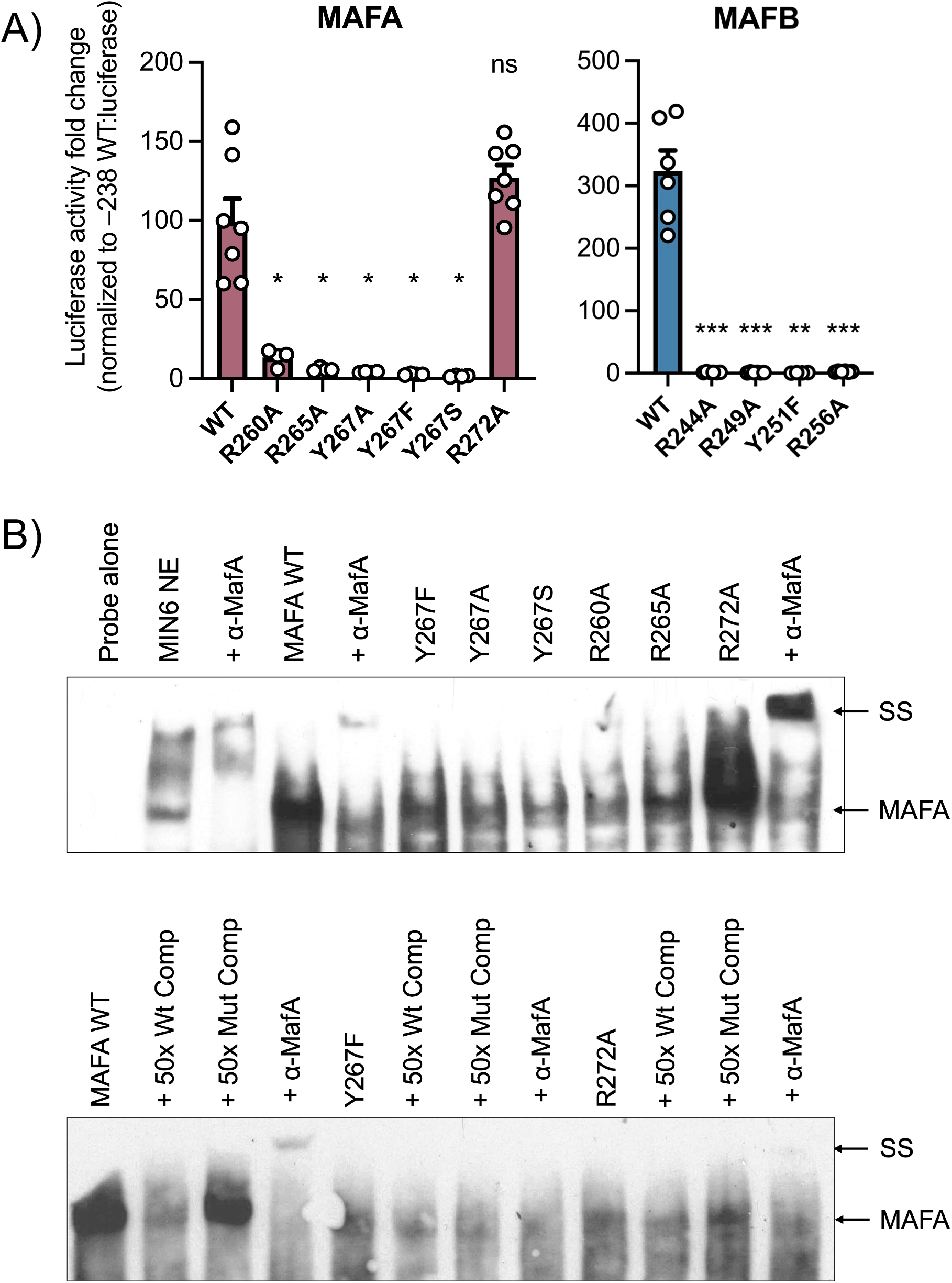
*Insulin* enhancer and DNA binding activity is retained in MAFA^R272A^, but not other conserved MAFA or MAFB basic regions identified in crystallographic DNA binding analysis. A) WT and mutant MAFA (left) and MAFB (right) expression vectors were transiently transfected into HeLa cells with the *Insulin* -238 WT reporter. All error bars indicate SEM. \*\*\**P*<0.001; \*\**P*<0.01; \**P*<0.05. n=3-4. B) Gel shift assays of HeLa nuclear extract produced WT and mutant MAFA. Top: Only MAFA^WT^ and MAFA^R272A^ were super-shifted (SS) upon addition of MAFA antibody. Bottom: MAFA^Y267F^ neither competed with WT versus mutant *Insulin* oligo nor super-shifted with MAFA antibody, whereas both MAFA^WT^ and MAFA^R272A^ did.

### 2) AlphaFold 2 predicts structurally unique regions within the MAFA, MAFB, and MAFA^S64F^ proteins

The level of sequence identity between MAFA and MAFB is variable between domains: the N-terminal *trans*-activation domain (51%), histidine-rich region (29%), extended homology region (66%), basic (100%), leucine-zipper dimerization region (63%), and C-terminus tail (8%) (Conrad et al., 2015) (**Figure 3A**). Structural information outside the basic-leucine-zipper region of MAFA and MAFB is likely of importance, as for example, preventing priming phosphorylation at S65 in MAFA and subsequently GSK3-mediated phosphorylation at serine (S) 61, threonine (T) 57, T53, and S49 unpredictably, and significantly changes SDS-PAGE mobility (i.e. 46 to 42 kD (Guo et al., 2009; Rocques et al., 2007). In addition, the MAFA DNA-binding ability, but not MAFB, appears to be regulated by phosphorylation within the N-terminal *trans*-activation domain region (Guo et al., 2010).

**Figure 3.**
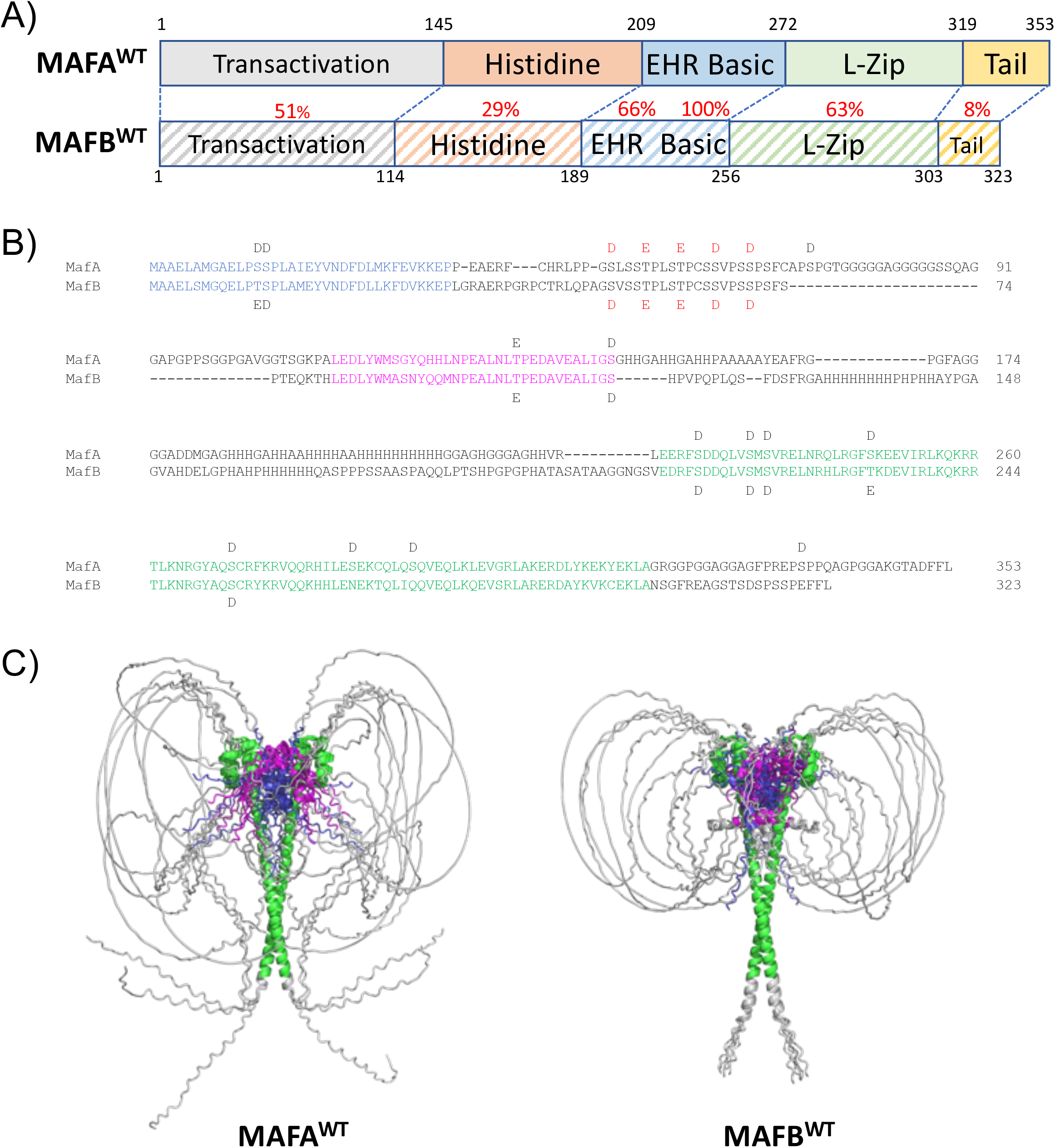
AlphaFold 2 modeling of MAFA^WT^ and MAFB^WT^ protein. A) Protein sequence identity (%) between various domains of MAFA^WT^ and MAFB^WT^ are denoted in red. Extended homology domain (EHR); Leucine-zipper (L-Zip). B) The aligned amino acid sequences of MAFA^WT^ and MAFB^WT^, with aspartic acid (D) and glutamic acid (E) used as phosphomimetics at *bona fide* sites of phosphorylation (Guo et al., 2010). The red labelled E and D represent the phosphosites absent in the underphosphorylated variants MAFA^S64F^ and MAFB^S70A^. The regions marked in blue (i.e., 83%) and magenta (i.e., 79%) have higher sequence conservation than the overall *trans*-activation domain and have structural elements in the AlphaFold 2 models. The region in green is the basic-leucine-zipper domain structured by AlphaFold 2 and determined by crystallography (i.e., MAFA residues 226–318 and MAFB residues 211–302) (Lu et al., 2012a; Textor et al., 2007). C) Comparing the five independently derived models for MAFA^WT^ (left) and MAFB^WT^ (right) illustrates potential structural difference in many areas of these proteins. For example, the longer C-terminal tail of MAFA leads to a more divergent conformation that can interact with N-terminal region sequences.

MAFA and MAFB act as homo- or heterodimers under physiological conditions (Tsuchiya et al., 2015). Here we employed AlphaFold 2 for *de novo* modeling of homodimers of full-length MAFA and MAFB. AlphaFold 2 employs deep iterative learning to provide predictions of the three-dimensional architecture of a protein solely based on multiple sequence alignments (Jumper et al., 2021). The analysis was performed on unphosphorylated MAFA (MAFA^un^) and MAFB (MAFB^un^), fully phosphorylated MAFA (MAFA^WT^) and MAFB (MAFB^WT^), and partially phosphorylated mutant MAFA (MAFA^S64F^) and MAFB (MAFB^S70A^). MAFA^un^ (and presumably MAFB^un^) is unable to bind DNA (Guo et al., 2010). For the purpose of modeling, aspartic (D) and glutamic (E) acid were used as phosphomimetics at S and T, respectively (**Figure 3B**). MAFA^S64F^ and MAFB^S70A^ have a reduced phosphorylation state relative to the WT protein due to the inability of a yet unidentified priming kinase to act at S65 in MAFA^S64F^ or S70 in MAFB^S70A^ (**Supp Figure 3A**).

Using a locally installed version of AlphaFold 2, we obtained five intrinsic structural models of the MAFA^WT^ and MAFB^WT^. This allows for a more robust exploration of conformational ensembles. The models for each of the sequences yields dimers with the expected folds within the basic-leucine-zipper domains, while the *trans*-activation domain had no specific conformation (**Figure 3C**). Of note are differences in the non-conserved C-terminal tail between MAFA and MAFB as well as the highly variable loops comprising the histidine-rich regions and the center of the *trans*-activation domain. The amino- and carboxy-termini of the *trans*-activation domain in the models display some secondary structure. Although there is no consensus on their absolute placement, they invariably lie near the DNA binding region of the basic-leucine-zipper domain.

Comparison of MAFA^WT^ to MAFA^un^ or MAFB^WT^ to MAFB^un^ indicates that the phosphomimetics result in a change of the AlphaFold 2 predicted models (**Supp Figure 3B, C**). Moreover, reducing the extent of phosphorylation in MAFA^S64F^ in comparison to MAFA^WT^ leads to further alterations in the placement of the *trans*-activation domain relative to the leucine zipper, which is depicted upon measuring the distances from Cα in each amino acid residue to a fixed point in space in the proteins (**Supp Figure 3D**). The pattern is slightly different in MAFB as the reduction in MAFB^S70A^ phosphorylation does not lead to a predicted change in the location of the *trans*-activation domain (**Supp Figure 3C, D**). Comparison of MAFA^WT^ to MAFB^WT^ indicates small differences in the regions proximal to the leucine-zipper domain and more substantial differences in the unstructured *trans*-activation domains. However, the most striking change between MAFA^WT^ and MAFA^S64F^ or between MAFB^WT^ and MAFB^S70A^ is in their C-terminal region (**Supp Figure 3B, C**), which is much more disordered in MAFA^WT^ and MAFA^S64F^ and in some models is found near the *trans*-activation domain. Next, we focused on determining how these possible structural differences relate to MAFA and MAFB activity.

### 3) WT and mutant MAFA and MAFB differ in their ability to functionally interact with other islet-enriched TFs

MAFA and MAFB, like many other TFs important in cell development and adult function, act in concert by binding and regulating enhancer activity (Scoville et al., 2015). These proteins bind to closely related *cis*-acting element sequences (**Figure 4A**) and are important in *Insulin* gene transcription in mice and humans (Artner et al., 2010), with, for example, MAFB being absolutely required for *Insulin* expression in human embryonic stem cell derived β-like cells (Russell et al., 2020).

**Figure 4.**
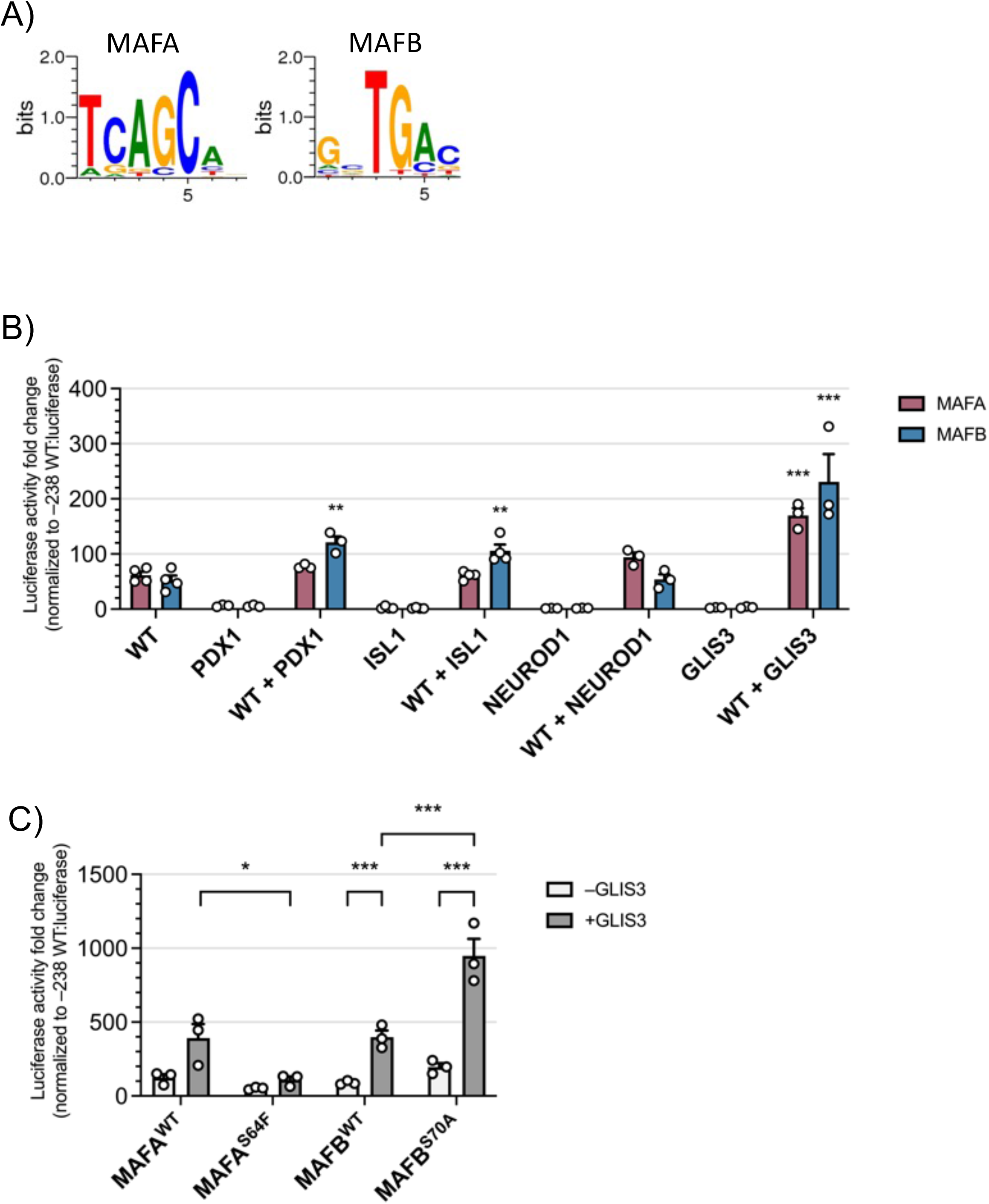
Comparing the ability of WT and mutant forms of MAFA and MAFB to act with other islet-enriched TFs to stimulate *Insulin*-driven reporter activity. A) The DNA binding sequence motif of MAFA and MAFB (Xie et al., 2009) as determined by ChIP-seq databases. B) Dual luciferase reporter assays were performed in HeLa cells transfected with vectors producing MAFA^WT^ (red bars), MAFB^WT^ (blue bars), PDX1, ISL1, NEUROD1, and/or GLIS3. The fold-change in activity of the -238 WT reporter over the no TF transfected controls is shown. Two-way ANOVA with Dunnett’s multiple comparisons test was performed. All error bars indicate SEM. \*\*\**P*<0.001; \*\**P*<0.01. n=3-4. C) Dual luciferase reporter assays were performed in HeLa cells transfected with vectors that express MAFA^WT^, MAFB^WT^, MAFA^S64F^, MAFB^S70A^ and/or GLIS3. The fold-change in activity of the -238 WT reporter over the no TF transfected control is shown. Two-way ANOVA with Tukey’s multiple comparisons test was performed. All error bars indicate SEM. ***P<0.001, **P<0.01; *P<0.05. n=4.

To determine if MAFA^WT^ and MAFB^WT^ differentially impact activation by other islet-enriched TFs of the *Insulin* gene, these proteins were co-transfected with PDX1, NEUROD1, ISL1, and GLIS3 and analyzed for their ability to increase *Insulin*-driven - 238 WT:luciferase reporter activity in HeLa cells. Notably, only the MAFA and MAFB proteins were able to independently stimulate *Insulin*-reporter activity (**Figure 4B**). MAFB, but not MAFA, provided additional enhanced activation in the presence of PDX1 and ISL1, while both MAFA and MAFB functioned cooperatively with GLIS3. In contrast, NEUROD1 did not elevate MAFA^WT^ or MAFB^WT^ activity, although the protein was expressed effectively (**Figure 4B, Supp Figure 4**). Differences were also observed in the ability of MAFA^S64F^ and MAFB^S70A^ to functionally interact with GLIS3, as MAFB^S70A^ provided additional stimulation over MAFB^WT^, whereas this enhancement was lost in MAFA^S64F^ (**Figure 4C**). Collectively, these results suggested that structural differences between the MAFA and MAFB proteins influence their ability to functionally interact with other islet TFs.

### 4) Analyzing how the predicted structurally unique regions of MAFA and MAFB affect ISL1 and PDX1 activation

Chimeric proteins between MAFA and MAFB were generated to evaluate how their N- and C-terminal regions influenced ISL1 and PDX1 activation of the *Insulin* -238 WT reporter. Importantly, the chimeric proteins retained the unique three-dimensional properties predicted by AlphaFold 2 in the corresponding domains of the WT protein (**Figure 5A and Supp Figure 5**). Interestingly, ISL1 stimulation was lost in MAFB^WT^ upon exchanging either MAFA^WT^ or MAFA^S64F^ N-(i.e., termed MAFA/B wherein amino acids 1-208 of MAFA were linked to residues 189-323 of MAFB^WT^) or C-terminal sequences (i.e., MAFB/A) (**Figure 5B**). These results indicated that the structural integrity of both the N- and C-terminal regions of MAFB were necessary for ISL1 stimulation.

**Figure 5.**
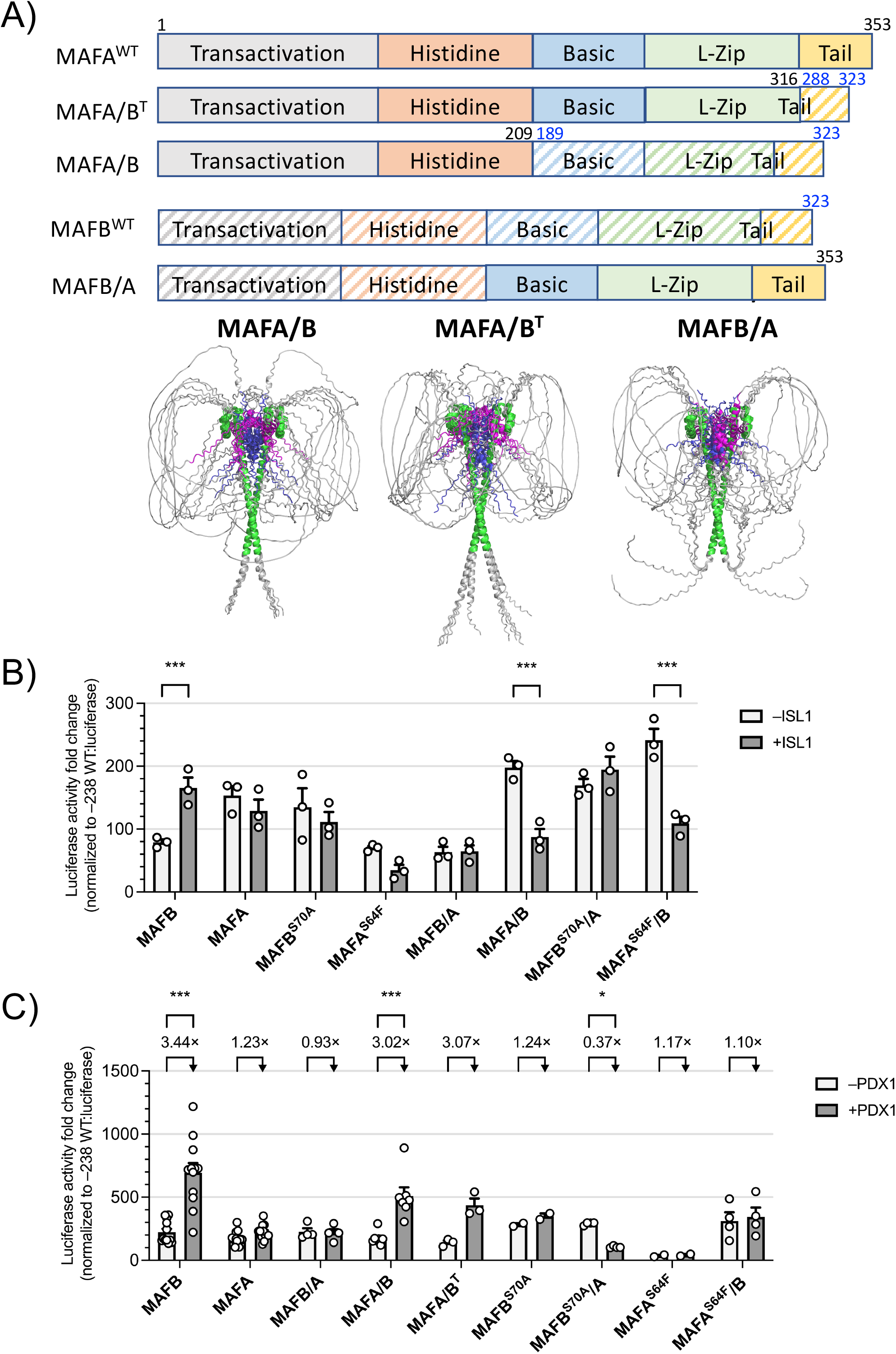
ISL1 does not activate the MAFA/B and MAFB/A chimeras whereas PDX1 activation requires MAFB C-terminal sequences. A) Top: The regions of the MAFA and MAFB chimeric proteins. Bottom: The models for the MAFA/B (left), MAFA/B^T^ (center), and MAFB/A (right) chimeras. B) Dual luciferase reporter assays were performed in HeLa cells transfected with vectors that express MAFB^WT^, MAFA^WT^, MAFB^S70A^, MAFA^S64F^, MAFB/A, MAFA/B, MAFB^S70A^/A, MAFA^S64F^/B, and/or ISL1. The fold-change in activity of the -238 WT reporter over the no TF transfected control is shown. Two-way ANOVA with Tukey’s multiple comparisons test was performed. All error bars indicate SEM. ***P<0.001, **P<0.01; *P<0.05. n=4. C) Dual luciferase reporter assays were performed in HeLa cells transfected with vectors that express MAFB^WT^, MAFA^WT^, MAFB/A, MAFA/B, MAFA/B^T^, MAFB^S70A^, MAFB^S70A^/A, MAFA^S64F^, and MAFA^S64F^/B with or without PDX1. The fold-change in activity of the -238 WT reporter over the no TF transfected control is shown. The relative fold changes of +PDX1 over -PDX1 were also shown. Two-way ANOVA with Turkey’s multiple comparisons test was performed. All error bars indicate SEM. ***P<0.001, *P<0.05. n=4.

In contrast to ISL1, MAFB^WT^ plus PDX1 activity was retained in N-terminal MAFA/B fusions (**Figure 5C**). Interestingly, the MAFA/B^T^ chimera that merely retained 35 C-terminal amino acids of MAFB^WT^ also appeared to work together with PDX1, although basal activity was reduced in the MAFB C-terminal fusions (i.e., MAFA/B^T^ by ∼84% and MAFA/B by ∼54%). Conversely, PDX1 plus MAFA/B activity was lost in MAFA^S64F^/B. In addition, stimulation with PDX1 was not observed in MAFB/A and even reduced by MAFB^S70A^/A. Collectively, these results imply that the distinct structural features within full-length MAFA and MAFB mediate key functional interactions with other essential islet-enriched TFs.

## DISCUSSION

MAFA^WT^ and MAFB^WT^ play crucial roles in a variety of cell types including pancreatic islet cells (Tsuchiya et al., 2015) across development and postnatal life. In this study, we first analyzed how mutants constructed within crystallographically determined basic region-DNA control element contact sites impacted protein activation. Notably, MAFA^R272A^ retained WT-like *trans*-activation and DNA-binding activity while the analogous MAFB^R256A^ mutant was inactive. These results implied that sequences residing outside the structurally analyzed basic-leucine-zipper region of MAFA and MAFB were functionally important. Consequently, the deep learning AlphaFold 2 algorithm was used to predict the three-dimensional architecture of the disordered regions within the non-DNA binding/dimerization regions of the WT and a MAFA or MAFB *trans*-activation domain mutant protein, and then analyzed for their potential functional impact. Predicted differences in the N-terminal and C-terminal disordered regions of MAFA^WT^, MAFA^S64F^ and MAFB^WT^ were found to regulate *Insulin* gene activation in cooperation with islet-enriched PDX1, ISL1, and GLIS3 TFs. Overall, we believe that the newly defined structural regions identified are essential to MAFA and MAFB activity.

The mutations identified within the basic region by X-ray crystallography would be expected to compromise TF DNA-binding ability. This expected property was found for all of the full-length MAFB mutants analyzed and for all but one in MAFA: MAFA^R272A^. It is not entirely clear why this discrepancy occurred in MAFA, but may reflect that the earlier structural analysis was performed on only a portion of an unmodified *Escherichia coli* expressed protein (i.e., aa 226-318 (Lu et al., 2012a)), appreciating that MAFA^WT^ and MAFB^WT^ are normally heavily modified in mammalian cells (e.g., phosphorylation (Guo et al., 2010), ubiquitination (Guo et al., 2009), and sumoylation (Shao and Cobb, 2009)). This likely includes the novel structural features detected by AlphaFold 2 in the full-length phosphomimetic substituted proteins (**Figure 3D**), presumably eliminating the expected role of MAFA^R272^ in DNA-binding.

The C-terminal tail structure of MAFA and MAFB predicted by AlphaFold 2 had the most striking impact experimentally. Thus, ISL1 activation by MAFB of the *Insulin*-driven reporter was prevented in the MAFB/A chimera (**Figure 5B**), while the MAFB^WT^ C-terminal conveyed PDX1 activation to MAFA/B and MAFA/B^T^ mutants and compromised basal MAFA^WT^ *trans*-activation activity (**Figure 5C**). Earlier studies had also revealed novel regulatory properties for the MAFA C-terminal tail. Hence, ectopic expression of MAFA, but not MAFB, promoted insulin^+^ cell formation within embryonic endoderm in chick *in ovo* electroporation assays (Artner et al., 2008). Moreover, only the chimeric MAFB/A protein induced chick *Insulin* activation and not a MAFA/B fusion (Artner et al., 2008). The inability of MAFB to stimulate *Insulin* expression in the *in ovo* electroporation assay implies that cooperating factors mediating activation during mouse (Artner et al., 2008) and human (Russell et al., 2020) pancreatic development are lacking in chick endoderm.

The analysis of PDX1, NEUROD1, ISL1, and GLIS3 on MAFA^WT^ and MAFB^WT^ activity was designed to glean whether their structurally distinct disordered regions predicted by AlphaFold 2 were regulatory. While it is presently unclear how much change DNA-binding would have imparted to these predictions, we believe that the functional interactions between PDX1, ISL1 and GLIS3 with the WT and chimeric proteins reveals their relevance to *Insulin*-driven expression. Consequently, we propose that collaboration between TFs like those analyzed here are important in regulating MAFA^WT^ and MAFB^WT^ controlled enhancers *in vivo*, many of which will be in common and some distinct. Our results demonstrated that such regulation entails recruitment of shared (e.g., GLIS3) and dissimilar (e.g., ISL1 by MAFB) TFs.

It is noteworthy that MAFA and MAFB are expressed in a distinct manner in mouse and human islet β cells, with only MAFA produced in mouse β cells and MAFA and MAFB in human β cells (Hang and Stein, 2011). In fact, mis-expression of MAFB is unable to rescue the many deficiencies associated with mice lacking MAFA in β cells (Cyphert et al., 2019). These TFs are not often co-expressed in human tissues, the exception being testes, islets, and skeletal muscle (**Supp Figure 6**). In these unusual circumstances, MAFA and MAFB presumably act in a heterodimer complex, which has been linked with the most active human islet β cell population (Shrestha et al., 2021).

Accordingly, the subpopulations of islet β cells expressing both MAFA and MAFB exhibit greater expression of key genes related to glucose sensing and hormone secretion as well as enhanced electrophysiological activity relative to subpopulations expressing only one or neither TF. Future efforts should be aimed at directly determining the structural and functional impact of the MAFA/MAFB heterodimer in this context.

## ACKNOWLEDGEMENTS

This research was performed using resources and/or funding provided by NIH grants to R.S. (R01 DK090570, R01 DK126482), J.C. (K08 DK132507), E.M.W. (F32 DK109577), and to the Vanderbilt Diabetes Research and Training Center (DK20593). J.C. was also supported by a Doris Duke Foundation Physician Scientist Fellowship (2020063) and a Burroughs Wellcome Fund Career Award for Medical Scientists, X.T. by a JDRF Fellowship (3-PDF-2019-738-A-N) and the Diabetes Research Connection (2022), and K.C.C. by a Career Development Award-2 (IK2BX006210) from the Veterans Health Administration. We thank the Center for Applied Artificial Intelligence in Protein Dynamics at Vanderbilt University for discussions related to the generation and interpretation of AlphaFold 2 predicted structures.

## MATERIALS AND METHODS

### Modeling by AlphaFold 2

Multiple sequence alignments were carried out with MMSeqs2 (Mirdita et al., 2019) and AlphaFold 2 models were generated with ColabFold (Jumper et al., 2021; Mirdita et al., 2022) using model parameters version 2.3. To allow for a more robust exploration of conformational space each alignment was run with 5 seeds leading to 25 models. To examine the effect of phosphorylation on the generated models, we used phosphomimetics based on previously identified phosphorylation sites in MAFA (Guo et al., 2009). Specifically, serine (S) residues were replaced with aspartic acid (D), and threonine (T) residues were replaced with glutamic acid (E, **Figure 3B**). These substitutions were made across the multiple sequence alignments excluding gaps. All of the phosphorylation sites in MAFA^WT^ and their homologous sites in MAFB^WT^ were replaced, whereas in conditions that mimic partial phosphorylation, all of the phosphorylation sites were replaced except for residues 49, 53, 57, 61, and 65 in MAFA^S64F^ or residues 54, 58, 62, and 66 in MAFB^S70A^. The highest predicted interface for each of the modelers is shown. To quantitate the variability in the AlphaFold 2 models, distances from Cα for each residue to a fixed point in space were measured. The fixed point was the centroid between R281 in MAFA, MAFA/B, and MAFA/B^T^, R259 in MAFB and R264 in MAFB/A. The distances were averaged for each set of 25 models.

### DNA constructs

The CMV-driven human MAFA and MAFB expression constructs used in the transfections were generated by Vector Builder Inc. (Chicago, IL). DNA sequencing analysis through Genewiz (South Plainfield, NJ) confirmed the fidelity of each construct.

### Cell culture and assays

HeLa monolayer cells were maintained in Dulbecco’s Modified Eagle Medium (Invitrogen) supplemented with 10% heat inactivated fetal bovine serum. CMV-driven MAFA, MAFB, PDX1, GLIS3, ISL1, and/or NEUROD1 were co-transfected with the rat II *Insulin* enhancer -238 WT:firefly luciferase plasmid (Artner et al., 2010), and the phRL-TK Renilla luciferase internal control plasmid. Reporter activity was evaluated 48 hours after transfection using a dual-luciferase assay according to the manufacturer’s protocol (Promega). Firefly luciferase measurements were normalized to the phRL-TK activity. Western blotting and gel shift reactions were performed as previously described (Cha et al., 2023) with 10 μg of nuclear extract and 200 fmol of the biotin-labeled double-stranded human *INS* MAFA/B binding site probe mixed either alone, with unlabeled competitor DNAs or MAFA antibody (Bethyl Laboratories, Product # A700-067) in a 20 μL reaction system (LightShift Chemiluminescent EMSA Kit, Thermo Fisher Scientific) containing 1x binding buffer, 2.5% glycerol, 5 mM MgCl, 50 ng/μL of poly(dI-dC) (Thermo Fisher Scientific), and 0.05% NP-40 (Sigma). The WT and mutant MAFA/B binding site sequences were described previously (Cha et al., 2023). The reactions were separated on a 6% precast DNA retardation gel in 0.5% Tris borate-EDTA buffer (TBE, Thermo Fisher Scientific) at 100 V for 1.5 hours. Each experiment was repeated at least three times using independently isolated plasmid preparations.

## Notes

### Competing Interest Statement

The authors have declared no competing interest.

### Summary of Updates

Major revision with addition of new data and updated Main figures (Figures 1-5); Authors updated; Supplemental files updated (Supplemental Figures 1-6)

