## Supplemental Figures for "Defining unique structural features in the MAFA and MAFB transcription factors that control *Insulin* gene activity"

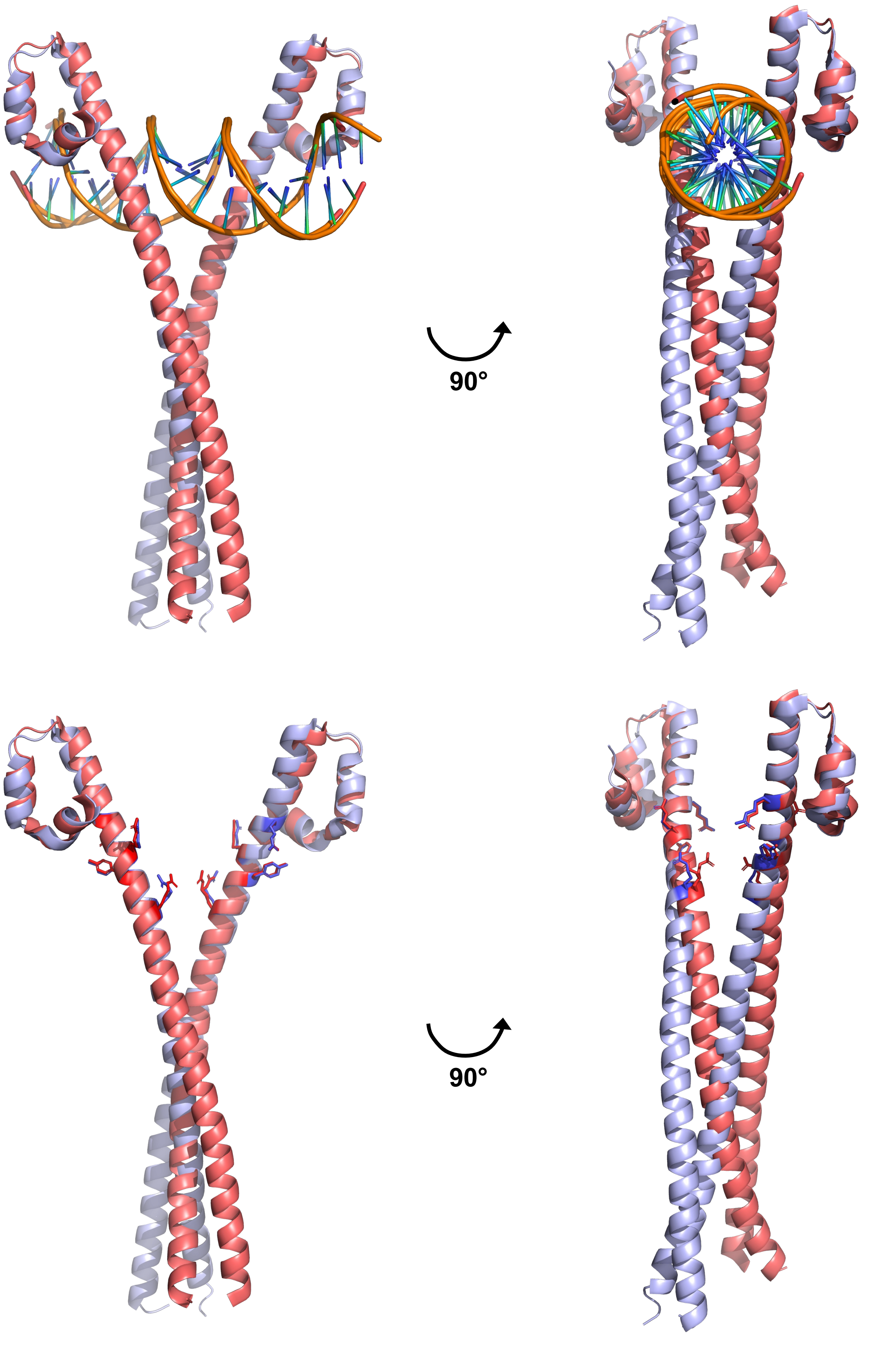


**Supplemental Figure 1**

A) MAFA

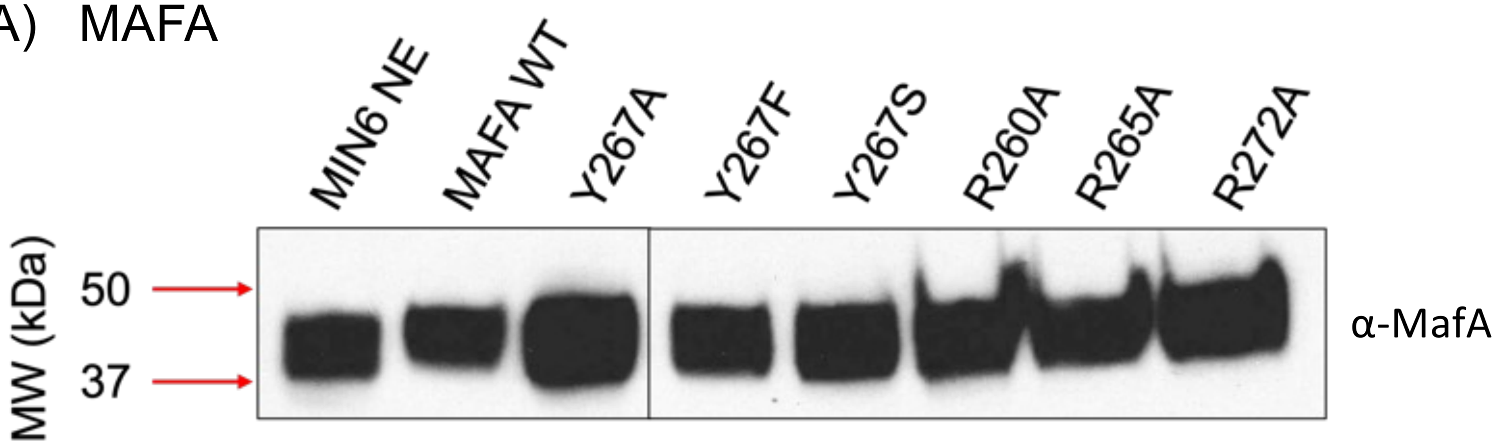


B) MAFB

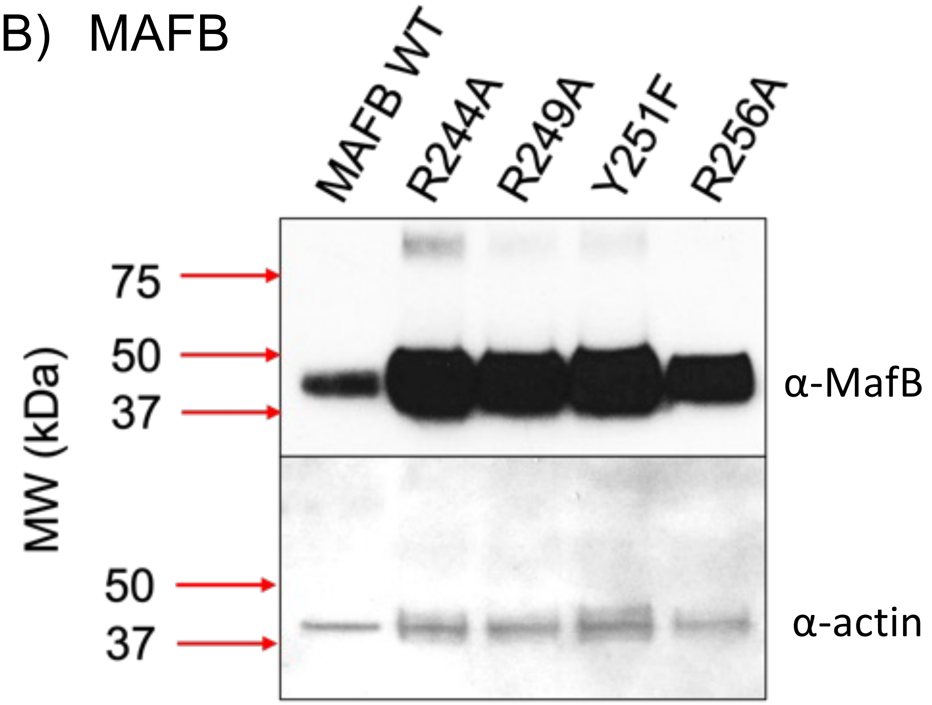


**Supplemental Figure 2**

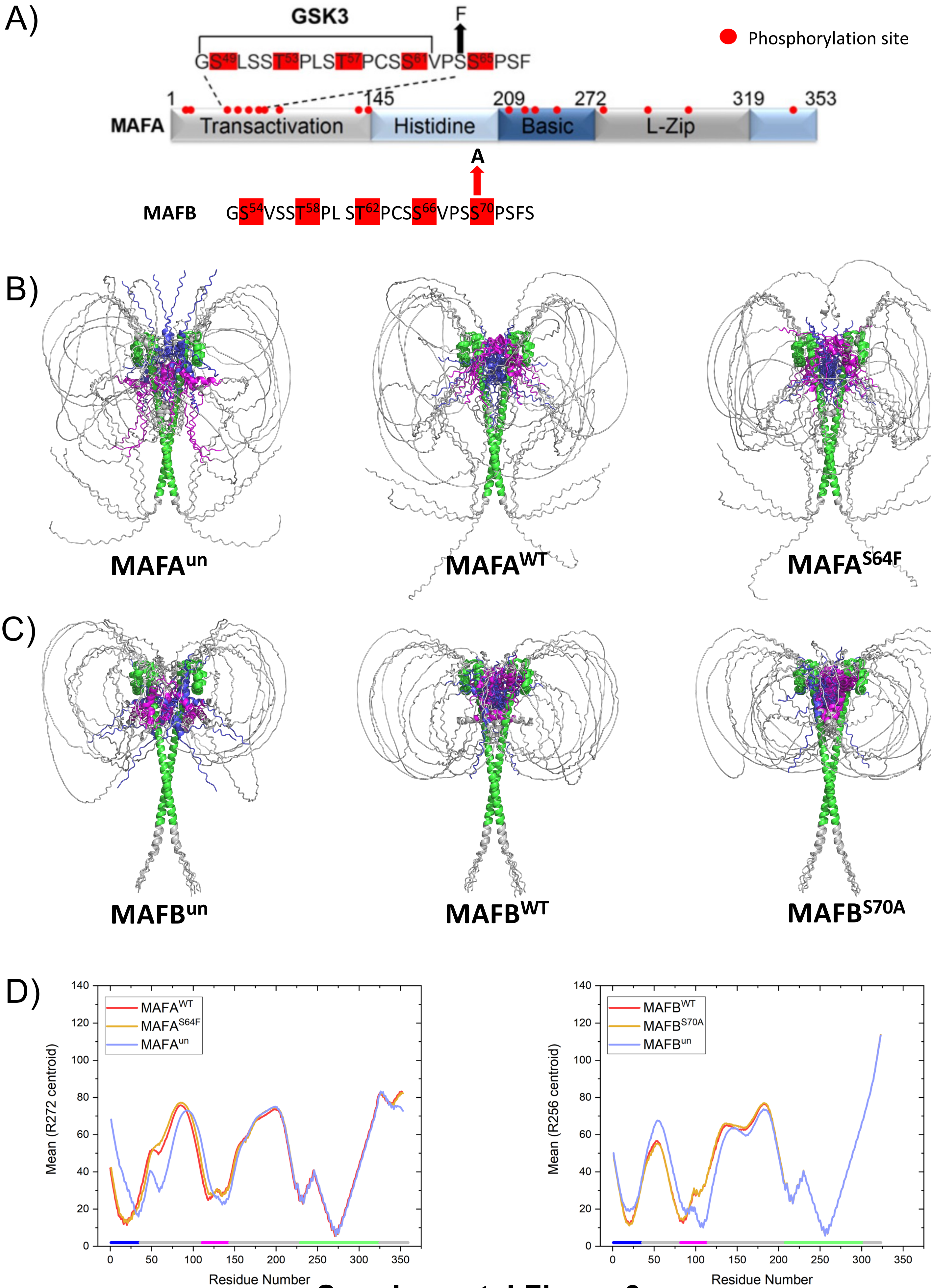


**Supplemental Figure 3**

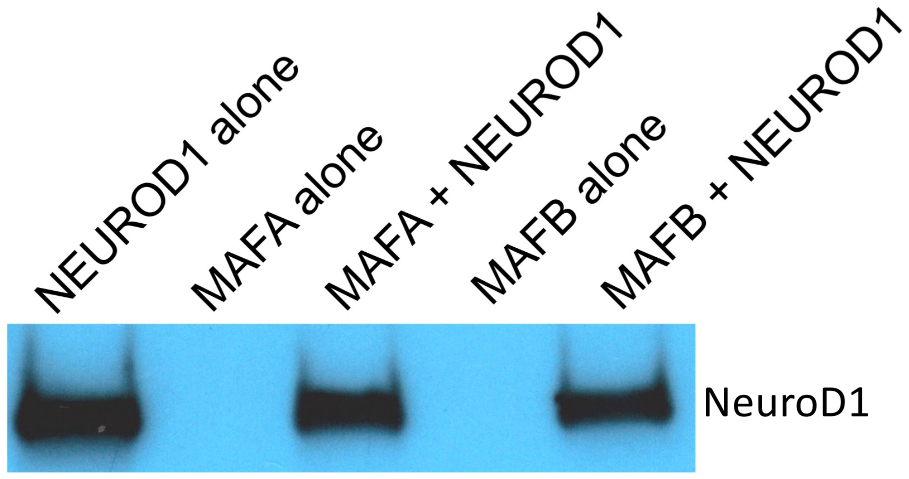


**Supplementary Figure 4**

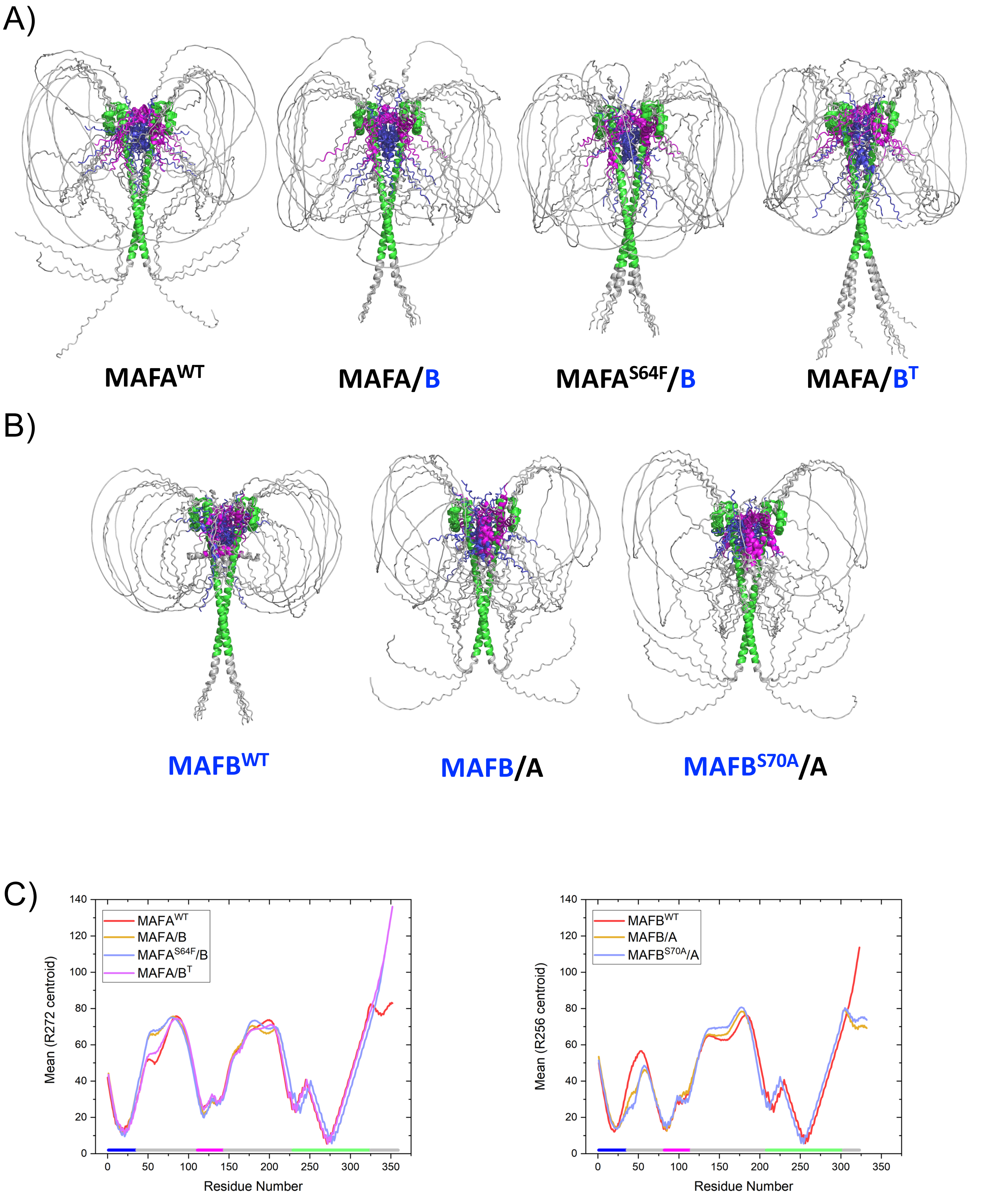


**Supplemental Figure 5**

A)

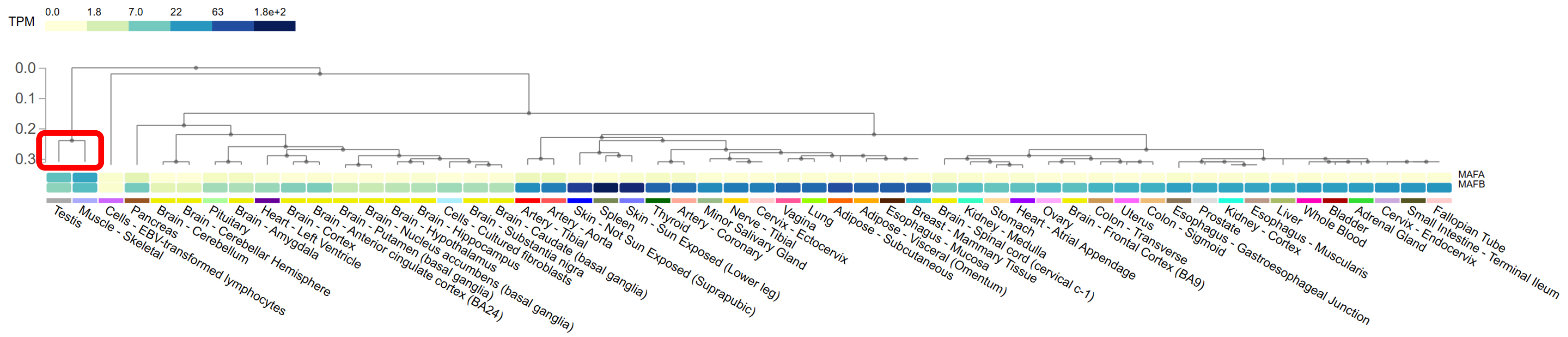


B)

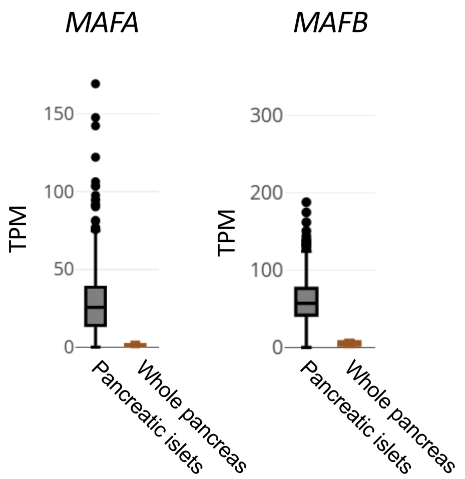


**Supplementary Figure 6**
